# On the prediction of arginine glycation using artificial neural networks

**DOI:** 10.1101/2022.06.05.494871

**Authors:** Ulices Que-Salinas, Dulce Martinez-Peon, A. D. Reyes-Figueroa, Ivonne Ibarra, Christian Quintus Scheckhuber

## Abstract

One of the hallmarks of diabetes is an increased modification of cellular proteins. The most prominent type of modification stems from the reaction of methylglyoxal with arginine and lysine residues, leading to structural and functional impairments of target proteins. For lysine glycation, several algorithms allow a prediction of occurrence, thus making it possible to pinpoint likely targets. However, according to our knowledge, no approaches have been published for predicting the likelihood of arginine glycation. There are indications that arginine and not lysine is the most prominent target for the toxic dialdehyde. One of the reasons why there is no arginine glycation predictor is the limited availability of quantitative data. Here we used a recently published high-quality dataset of arginine modification probabilities to employ an artificial neural network strategy. Despite the limited data availability, our results achieve an accuracy of about 75% of correctly predicting the exact value of the glycation probability of an arginine-containing peptide without setting thresholds upon whether it is decided if a given arginine is modified or not. This contribution suggests a possible solution for predicting arginine glycation. Our approach will greatly aid researchers in narrowing down possible glycation sites in protein targets. This strategy could improve the structural and functional characterization of proteins of interest.

## INTRODUCTION

In nature, there is an amazing variety of proteins. So far, thousands of them have been described that carry out diverse functions that are essential for life, either as structural building blocks within and without cells or as catalysts of biochemical reactions in the form of enzymes ^1,2^. Proteins are composed of a certain number of amino acids. There are twenty ‘standard’ amino acids and several somewhat obscure ones like selenocysteine and pyrrolysine that are also proteinogenic.

Clearly, the potential sequence variety is enormous even in short proteins (peptides), reaching astronomical proportions. To this huge variety, so-called post-translational modifications of amino acids must be added. These modifications add layers of regulation and control and include a plethora of processes leading to adducts, like acetylation, phosphorylation, methylation and ubiquitination among many others ^3^. The post-translational modification of specific amino acids can occur enzymatically or non-enzymatically. For example, protein glycosylation, which is important for protein sorting, protein secretion and cellular recognition among other functions, is performed by glycosyltransferases and related enzymes ^4^. Another important example is the reversible modification of histones by histone acetylases and deacetylases that is essential for the coordinated regulation of gene expression ^5^. Glycation, on the other hand, is regarded as a strictly non-enzymatic process that involves the reaction of sugars (e. g., glucose, fructose) and sugar-derived molecules with amino groups of biologically highly relevant molecules, like nucleic acids, lipids and proteins ^6^. Usually, these reactions result in the formation of advanced glycation end-products (AGEs) which are mostly detrimental and compromise the function of the target molecule irreversibly ^7,8^.

In proteins, the side chains of lysine and arginine are the main targets of AGE formation ^9,10^. One of the most reactive glycating compounds is the reactive carbonyl species (RCS) methylglyoxal (MGO) which is formed as a toxic by-product by metabolic activity, e. g. during glycolysis ^11^.

Usually, cellular MGO levels are kept at relatively low levels of around 0.3 to 6 μM ^12^ by a dedicated enzymatic defense systems (e.g., glyoxalase I and II, aldose reductases) ^13,14^ and low-molecular weight scavengers, but in certain pathological conditions (i. e., diabetes, neurodegeneration, cancer) ^15,16^ and in aged cells and tissues ^17–19^ MGO can become problematic for cellular viability due to increased production and/or impaired removal. It should be noted that the specific MGO-mediated modification of proteins can be important for several signaling processes and for gene regulation. This has been demonstrated in studies often conducted in simple eukaryotic model systems that are very amenable to experimental procedures ^20^.

Although the importance of MGO binding to certain amino acids in a target protein is a well-studied phenomenon, it has become clear that there are most likely no straightforward consensus sequences that allow a reliable prediction of potential glycation sites ^21^. Available predictive algorithms therefore must rely on the physical (e. g., polarity), chemical (e. g., amino acid composition) or structural features (e. g., accessible surface area, secondary structure features and local backbone angles) of nearby amino acid residues. These allow a prediction of potential sites of lysine glycation. For example, *GlyNN* ^22^ utilizes an artificial neural network (ANN) ^23^ approach to enable lysine glycation prediction from a relatively small dataset of 215 elements. Further developments are *BPB_GlySite* ^24^, *PreGly* ^25^, *PredGly* ^26^, *Gly-PseAAC* ^27^, *Glypre* ^28^, *iProtGly-SS* ^29^ and *GlyStruct* ^30^ with approaches like bi-profile Bayes feature extraction, position specific amino acid propensity and models trained with support vector machine (SVM) classifiers. Traditionally, lysine glycation has been the target of in-depth research with a large database of lysine modifications being freely available (PLMD ^31^, which is based on CPLM ^32^ and CPLA 1.0 ^33^). However, in recent years it has become clear that also, possibly even more so, the reaction of MGO and arginine is very relevant for the pathogenicity of AGEs ^9,34^. Chemically, MGO reacts with arginine yielding an irreversible intermediate, the AGE dihydroxyimidazoline (DHI) after Schiff base addition and its subsequent rearrangement (Amadori product formation). Removal of water from DHI leads to the formation of the AGE 5-hydro-5-methylimidazolone (MG-H1) ^7,8^. MG-H1 is an important marker for the AGE-modification of skin collagen ^35^, mitochondrial dysfunction ^36^ and acute coronary syndrome ^37^ among others.

The experimental demonstration of specific amino acid modifications is often very costly in terms of time, resources, and labor, especially in larger proteins which may contain several potential sites of glycation. Hence, being able to analyze protein sequences for the presence of glycated arginine residues would be a useful approach to predict sites of MG-H1 formation. To our knowledge, no tools have been published so far that allow the predictive identification of potential arginine modification sites in proteins. In this work, we implemented a machine learning (ML) method using our own supervised numerical training algorithm, which uses the features of amino acids as input information, and as target the probability of glycation to occur in a certain protein, where the selected amino acids form a sequence of eleven elements within the protein whose central amino acid will always be arginine. Specifically, we utilize Artificial Neural Networks (ANN) because they offer a direct way to solve problems given their high accuracy and their adaptation to noisy, unknown or incomplete information ^23^ besides a fast computation after training due the fact that the neural network can be easily implemented in the parallel hardware. In particular, ANNs as ML methods have been applied in problems of fluids in the flow phase pattern identification, for example ^38,39^.

The peptide sequences we used were extracted from Scheck *et al*. ^21^. Although the number of glycated peptides is relatively small for training or prediction, the information is of exceptional quality because all experiments were performed by the same lab using the same procedures for modification and its detection. As such our work is not based on glycation data from different sources that are inherently difficult to compare. Furthermore, our approach allows stating a probability of glycation in percent. It is therefore superior to algorithms that are based on thresholds that decide whether a residue is glycated or not. In conclusion, our contribution is aimed at enabling a more directed protein-AGE analysis saving time and funds for the researcher.

## METHODOLOGY

### GENERAL OUTLINE

We chose the following features of amino acids to computationally characterize a short protein sequence (11 amino acids) that contains a central arginine residue: sequence of amino acids of the peptide (SoA), hydropathy (Hyd), mass (Mas), hydrophobicity (Hyp), polarizability (Pol), normalized van der Waals volume (vdW), torsion angle (ToA), and isoelectric point (IEP) (Fig. 1). We subsequently formed vectors of the type ∈ R^1X11^ for each feature and selected the number of vectors, one (study case 1), two (study case 2), or three (study case 3), that enter an artificial neural network (ANN) to train a model that allows us to predict the glycation probability of the central arginine residue in the 11-mer peptide sequence. These sequences were taken from a recent publication by Scheck *et al*. ^21^. The extent of arginine modification was determined by these authors using a state-of-the-art technique (liquid chromatography mass spectrometry). In total, 54 sequences were retrieved. Although the number of glycated peptide fragments is not particularly high, all experimental steps were conducted under comparable conditions, making comparisons much more reliable ^21^. Fig. 1 shows a general outline of the process to be followed in our methodology. More details on the ANN operation can be found in the supplementary materials to this manuscript.

**Fig. 1:**
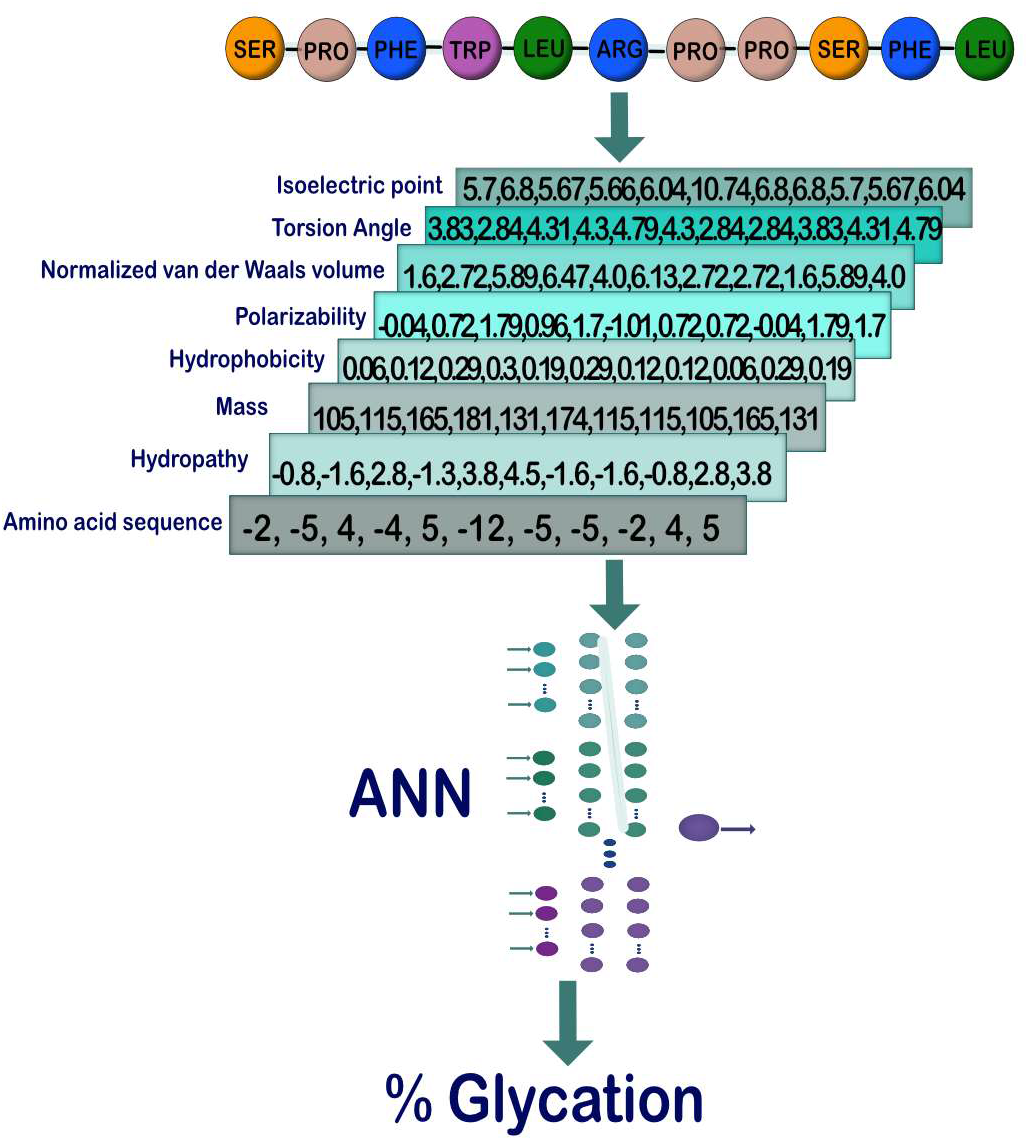
Steps to follow for the characterization and prediction of glycation using ANN. First, a preliminary database is assembled from the amino acid sequence of the peptides. Then, by rewriting the amino acid sequence with the values corresponding to each of the physical properties, a list of all the vectors is created. Their values are normalized and delivered to the ANN, which through a learning process makes the final predictions corresponding to the probability of glycation for each peptide.

### DATABASE CONSTRUCTION AND STUDY CASES

The tool of ML used to make the prediction of peptide glycation is an artificial neural network (ANN) that requires data for training and testing. As stated above, for the construction of the arginine glycation database, information obtained experimentally by Scheck *et al*. ^21^ was used. This information allows us to rewrite the alphabetical sequence of amino acids for each of the peptides with the corresponding numerical values for each of their physical properties. The list of the 20 proteinaceous amino acids, with their corresponding values for each of the physical properties that represent them can be found in Table S1 in the supplementary material.

By employing this information, we built a database with 54 (peptides) × 8 (features) vectors using the different values for each amino acid (11 elements). That is, each one of the 432 vectors was formed by selecting one of the 54 peptides made up of a sequence of 11 elements, and selecting one of the eight physical properties, thus assigning to each element of the peptide the value corresponding to that physical property. For example, for the 11mer-sequence SPFYLRPPSFL we built eight vectors (Fig. 2). We retrieved each element of the sequence (i. e., an amino acid) and obtained the corresponding values (for the complete list of constructed vectors refer to Tab. S2). This process was repeated for all properties.

**Fig. 2:**
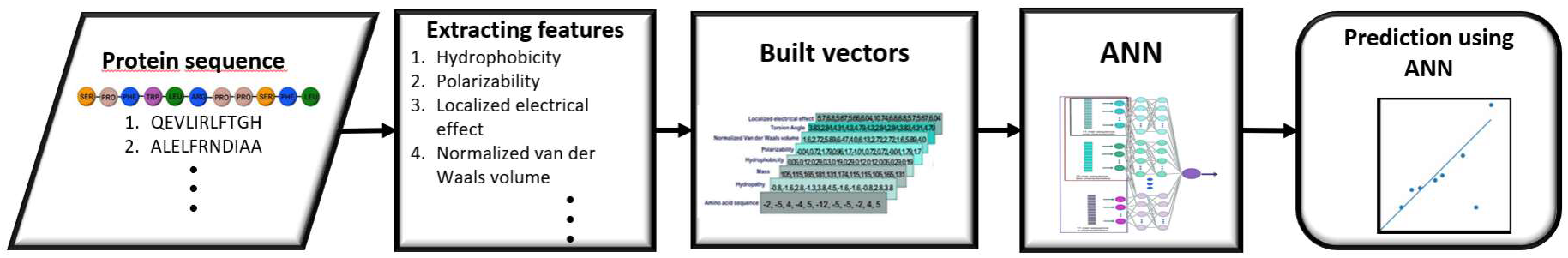
Example of construction of vectors. For the 11-sequence SPFYLRPPSFL, each amino acid is converted into a number, dependably of the property. The first amino acid, serine (SER), has a value of 5.7 for the property localized electrical effect, 3.83 for torsion angle, 1.6 for normalized van der Waals volume, -0.04 for polarizability, 0.06 for hydrophobicity, 105 of mass, -0.8 for hydropathy, and -2 for amino acid sequence.

We considered basically three cases as inputs for the ANN. The single-case study is accepting one individual vector of the same property for each peptide as input. The two-case study considers two vectors of two different properties in combination for each peptide, giving rise to up to 28 different outputs. Finally, the three-case study considers three vectors of three different properties in combination for each peptide, giving rise to up to 56 different outputs. We would like to point out that, considering that higher order combinations result in more complex learning processes without providing significant improvements in predictions. Consequently, we did not consider them further in our study.

Here it is important to note that, for any of the three cases, at the center of the sequence of elements is always the amino acid arginine. Recall that the objective of ANN in this project is to be able to predict the glycation probability of the central arginine corresponding to each peptide. This can be done through the combination of vectors as described above.

At this point it is worth taking a few steps forward and establishing now that another part of the objective of this study is to find out which of all these possible combinations of amino acids parameters gives us the most accurate predictions.

For the ANN learning and prediction process, it is necessary to form a set of samples for each of the study cases, which we termed patterns. Each pattern was built on a combination of only one, two or three vectors for a specific peptide where their numerical values are the properties used in the corresponding amino acid sequence of the peptide. Thus, each pattern (𝒫) is represented by a matrix of “m” rows (the number of properties selected) and “n” columns (each one of the eleven amino acids within the peptide).

These patterns are provided to the ANN to be able to predict the exact probability of glycation of the central amino acid arginine (inside the pattern).

For example, if we wanted to predict the probability of glycation of a certain peptide from the analysis of a pattern formed by the combination of two vectors corresponding to hydropathy (Hyd) and mass (Mas), we would specify an array *A*_*mn*_ of the form:

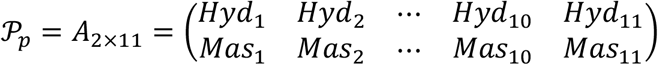

where p is the index for each one of the 54 peptides. Thus, for the 11-sequence SPFYLRPPSFL mentioned above, we can consider that for a combination of two vectors, to form a 2-vector pattern (𝒫), we can take any two of the 8 different sequences of numerical values shown in Fig 2.

All the specifications of the study cases are presented in Table 1 for reference. From the eight amino acid features, there are multiple combinations to conform each one of the study cases. For the two-case and three-case we will focus on the combinations of features that present the best results. However, for the one-case, to explain in detail, the learning and prediction process of the ANN, we have chosen to present the results of all the features, thus forming a total of eight sub-cases (Table 1).

**Table 1:** Details of the ANN study cases. The physical properties and the architecture of the neural network are presented for each one of the sub-cases of the case 1, along with the sub-cases that performed the best result for cases 2 and 3.

| Cases | Features | ANN layer architecture |
| --- | --- | --- |
| Case 1A | Sequence of Amino Acid | 11x4x3x2x1 |
| Case 1B | Hydropathy | 11x4x3x2x1 |
| Case 1C | Mass | 11x4x3x2x1 |
| Case 1D | Polarizability | 11x4x3x2x1 |
| Case 1E | Hydrophobicity | 11x4x3x2x1 |
| Case 1F | Normalized van der Waals volume | 11x4x3x2x1 |
| Case 1G | Torsion angle | 11x4x3x2x1 |
| Case 1H | Isoelectric point | 11x4x3x2x1 |
| Case 2 | Polarization + van der Waals volume | 22x10x1 |
| Case 3 | Sequence of Amino Acid + Hydropathy + van der Waals volume | 33x40x1 |

Taking up the fact that for each of the patterns used as input information for the ANN training is composed of an array of “m” features and eleven values the amino acids (n), then, for the three-case study, we will have an array of the form:

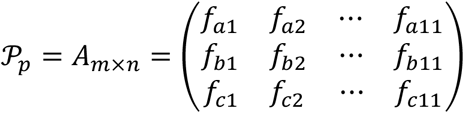

where the index p represents the peptide and therefore runs from 1 to 54, and the index m represents each one of the three features chosen (f_a_, f_b_ or f_c_) among the eight possible ones; where indices a, b and c take different values from each other, ranging from 1 to 8.

For optimal ANN performance, the input data for all cases are preprocessed to result in the normalized 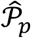 pattern, for which

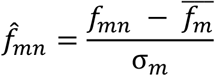

each feature m and for each amino acid n we will normalize the matrix elements following to the relation (3), where 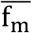 is the mean of the elements of the training set corresponding to the m-th feature, and σ_m_ is the standard deviation of those elements. The normalization process for a given case has been performed on each subset of vectors made up of the elements corresponding to the same feature, and not on the whole dataset.

For the learning process, all the normalized information will be divided into three sets, the first and largest will be used for training, consisting of 70% of the data. The second is the validation set consisting of 15% of the data and the remaining 15% will be used for the final predictions that will be presented in the results section. It is important to note that during training the ANN does not know the data of the validation and prediction sets.

Thus, the ANN will be fed only with the training set for each of the case studies. Where, in general, the ANN architecture is of one to three hidden layers, having per layer (including the input layer), a varying number of neurons, according to each of the cases studied (see Table 1).

Fig. 3 shows the general architecture of the ANN used. Consider that it will be fed with the patterns formed for each of the case studies, through the input layer of the ANN. Subsequently, learning is performed through the hidden layers and the prediction is processed in the output layer.

**Fig. 3:**
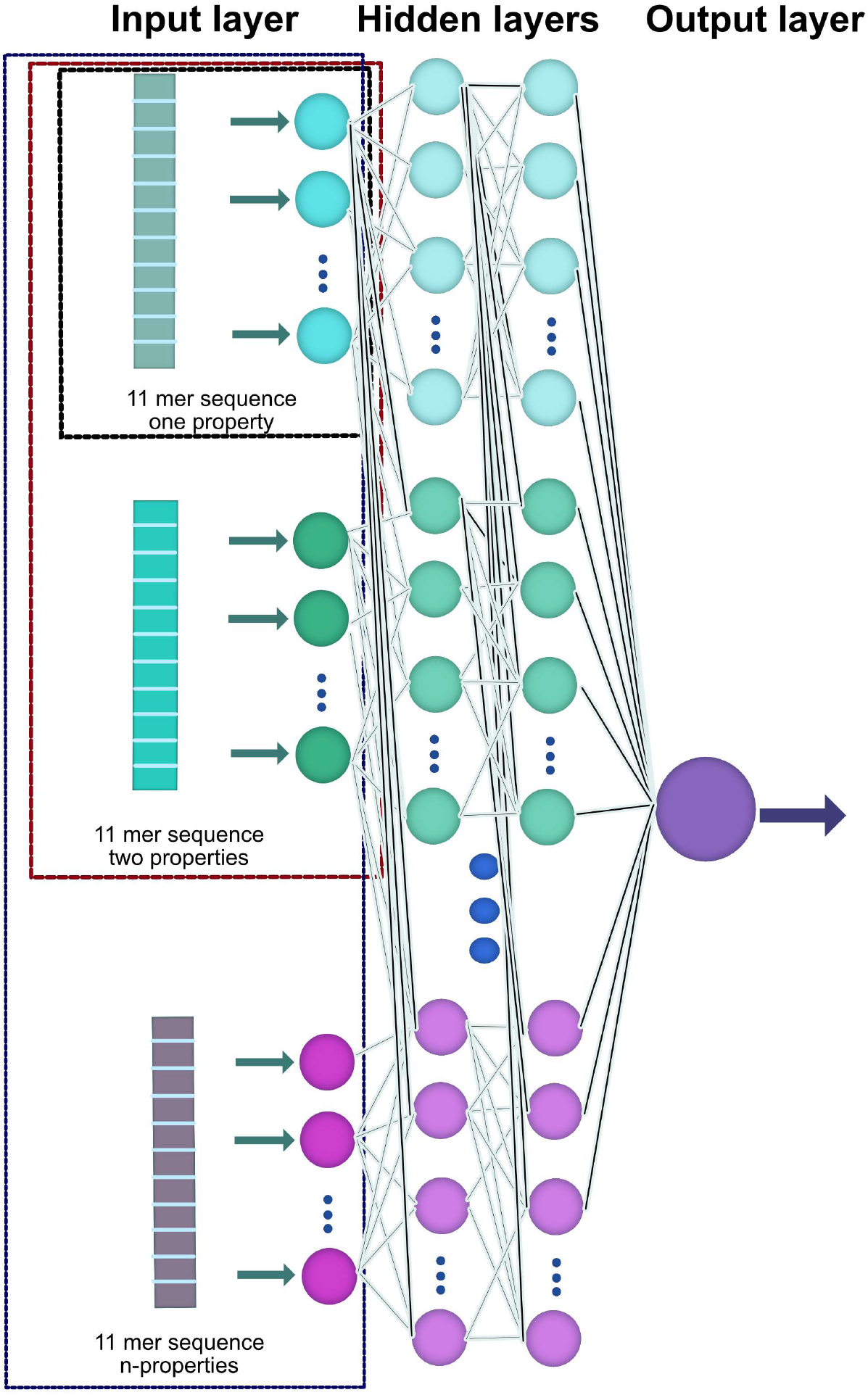
Schematic of the utilized ANN structure. Each of the 11mer sequences with n-features is provided to the ANN through the input layer, from which the learning process proceeds through an adjustment in the interconnections in the hidden layers. Finally, a prediction of the glycation probability is made, which is provided by the output layer.

Finally, to minimize the error during the training process the ANN was constructed as a regression model using the Adam optimization algorithm ^40^ with a learning rate γ=0.001, we have used a rectified linear unit (ReLu) as an activation function ^41^, and employing a backpropagation algorithm ^42^.

## RESULTS

Towards predicting the value of the glycation probability based on the small database available, ANNs were used. They can offer high accuracy, even with incomplete information ^23^. As previously described, we used eight different features to construct the vectors, feeding the algorithms with one of the features for each sequence, or using a combination of them. We report the Mean Absolute Percent Error (MAPE) and Mean Absolute Error (MAE), by averaging the results over 160 different predictions, each with a different ANN training.

Table 2 summarizes the analysis carried out from the eight different features and the combination of each two of them. The MAPE and MAE values are reported in the upper and lower diagonal matrices respectively. In the main diagonal of the matrix, the MAPE is represented first, followed by MAE. Both errors are also characterized by gray shades, to describe if we have a high accuracy (clear gray), average (medium gray) or low accuracy (dark gray) prediction value.

**Table 2:** Summary of glycation probability errors (MAPE/MAE) for the single-case / two-case predictive approaches. We report the MAPE values in the upper diagonal matrix, and MAE values in the lower diagonal. The main diagonal shows the MAPE and MAE value for the case 1 respectively. Both errors are also characterized by gray shades, to describe if we have a high accuracy (clear gray), average (medium gray) or low accuracy (dark gray) prediction value.

|  | MAPE |  |  |  |  |  |  |  |
| --- | --- | --- | --- | --- | --- | --- | --- | --- |
|  | *vdw* | *Mass* | *Pol* | *Ami* | *Hid* | *IdP* | *ToA* | *Hih* |
| *vdw* | 39.75/27.49 | 35.71 | 33.3 | 37.61 | 35.77 | 37.34 | 40.62 | 39.45 |
| *Mass* | 24.83 | 40.93/27.5 | 36.74 | 46.28 | 40 | 39.05 | 46.47 | 39.36 |
| *Pol* | 23.53 | 25.5 | 41.06/28.3 | 36.79 | 35.66 | 37.88 | 41.51 | 41.73 |
| *Ami* | 25.63 | 29.93 | 25.74 | 41.71/26.98 | 40 | 46.59 | 44.96 | 44.74 |
| *Hid* | 25.69 | 29.87 | 25.66 | 26.36 | 42.6/28.17 | 45.4 | 46.58 | 46.22 |
| *IdP* | 23.96 | 26.11 | 23.87 | 26.81 | 27.02 | 48.34/28.95 | 46.15 | 52.23 |
| *ToA* | 24.69 | 28.85 | 25.15 | 28.42 | 28.83 | 26.08 | 52.12/32.11 | 57.13 |
| *Hih* | 26.21 | 26.77 | 26.51 | 27.48 | 27.8 | 27.57 | 30.61 | 54.79/30.63 |

The cases where we use only one feature can be seen in the main diagonal, for MAE values the lower results we had were using Hyp (30.63) or ToA (32.11), whereas, we have the highest values using SoA (26.98) or vdW (27.49). Also, for MAPE the lower results we had again were using Hyp (54.79) or ToA (52.12), and for the highest results we have the best values using vdW (39.25) or Mass (40.93).

We have seen that the combination of features can improve the performance of the ANN, now we will review the results for each specific case, recall that the previous values were the average over 160 different predictions. Fig. 4 shows the box plots of MAPE and MAE for the individual features, even though vdW results in the lowest errors in the table 2, we can see a broader distribution, in contrast with Hyd. As we can see from table 2, Hyd represents a 7.2 and 2.5% higher MAPE and MAE error than vdW, but 66.7 and 70.7% narrower range distribution. The narrowest range distribution for MAPE and MAE are represented by IEP and Hyp respectively, in contrast, the broadest is Pol for both cases.

**Fig. 4:**
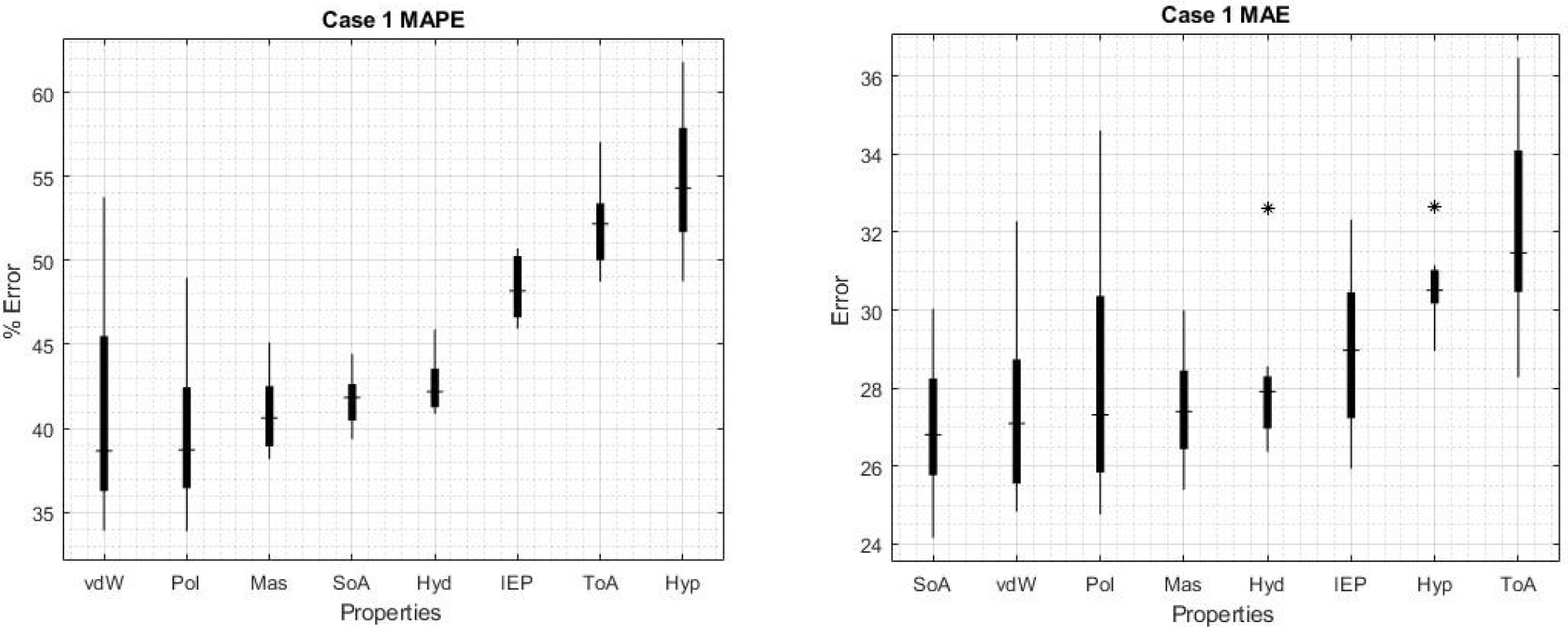
Box plot of the results for case 1. Values obtained for MAPE (left) and for MAE (right). The characteristics are ordered from the lowest error (implying higher reliability and accuracy) to the highest errors.

For cases 2 and 3, where there are multiple combinations of amino acid features, in a search for clarity it was considered to present in detail for this work, only the cases that showed the best results (considering that the remaining details are shown in the supplementary material, Fig. S1).

Thus, for case 2, the highest errors were obtained with the combination of Hyp and ToA with values for MAE and MAPE of 30.61 and 57.13, respectively. Now, if we use the best features in the main diagonal to make the combinations, we can improve the results compared to one feature only. The best value we were able to find using a combination of two features was vdW-Pol for both MAE and MAPE with values of 23.53 and 33.3 respectively (bold numbers in Table 2).

Fig. 5 shows the MAPE and MAE respectively of the first 8 combinations of two features with the lowest values. In turn, for comparison, the two combinations with the highest errors are presented (the complete set can be found in the supplementary materials, Fig. S1). Interestingly, once we study the combination of two features, most of the cases show error distributions as narrow as using only the feature Hyd, which is the single feature that shows the narrowest error distribution (see Fig 4). This can lead us to the idea that, increasing the number of features used in the ANN increases the performances and narrows the error distribution, but we must be aware that this is not always the case, because the features may hide unknown correlations, whereby increasing the number of them will not improve the prediction, as the amount of independent data may not raise. The narrowest error for case 2 is presented in the combination of Mas-ToA for MAPE and Hyd-IEP for MAE with a range of 2.13 and 0.80 respectively.

**Fig. 5:**
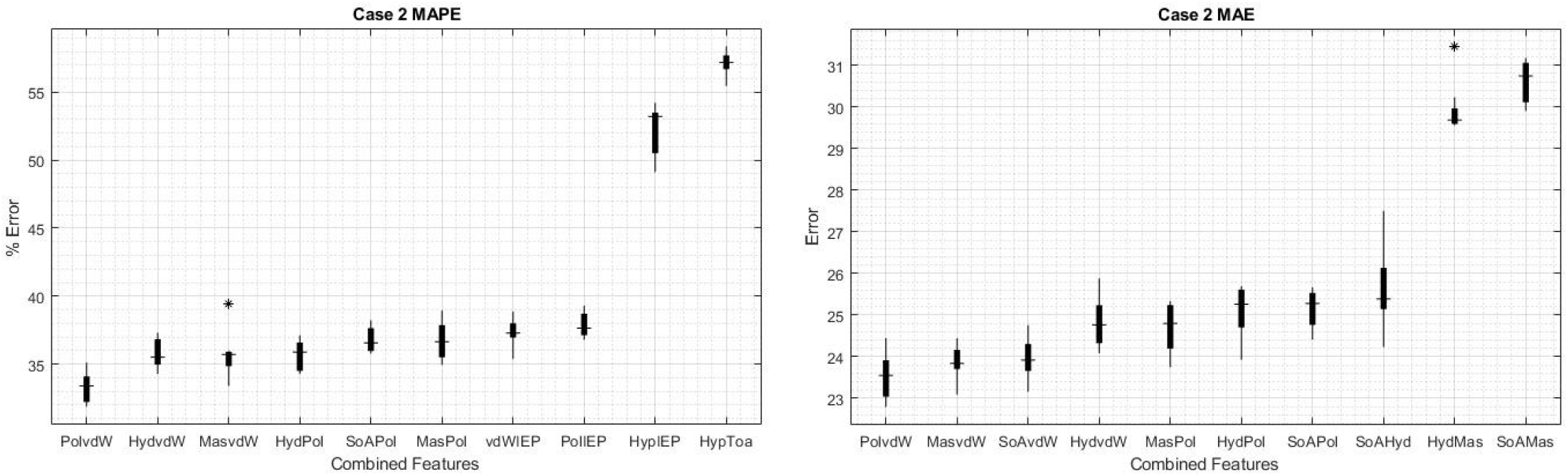
Box plots of the results for case 2. Values obtained for MAPE (left) and for MAE (right). The first eight combinations with the lowest errors are shown, as well as the two combinations with the highest errors.

In case 3, consisting of the combination of three different properties, an improvement in ANN performance was generally observed, mainly in combinations that incorporated the properties SoA, Hyd, Mas, vdW and IEP, whose trend can be seen from case 2 in Fig. 5. Thus, the lowest value for MAE was obtained by combining SoA with Mas and IEP, reaching an error of 15.42; while for MAPE a value of 25.04% was reached by combining Hyd with Mas and IEP; which is a substantial reduction with respect to the best results of case 2.

## DISCUSSION

Here we developed a tool for predicting the glycation probability of arginine residues in proteins. Although we had to work with a limited dataset, we consider it relevant to have some means to predict arginine modifications. Furthermore, our work can serve as a work of principle and be subsequently expanded once more information on arginine glycation becomes available.

Arginine, akin to lysine, is a prime target for methylglyoxal ^9^. Due to the fact our data set on arginine glycation is too small for conventional approaches, we employed a machine learning strategy. While artificial intelligence (AI) is the overarching science of mimicking human abilities, machine learning is a specific subset of AI that trains a machine how to learn. Nowadays, machine learning is one of the most important tools for scientists in the development of new applications ^43,44^. We could have conveniently employed a linear regression-based approach to estimate the probability of glycation, but the results would have been considerably poor compared to those obtained using the more sophisticated ANN.

At present, the accuracy of our algorithm is limited by the relatively small size of the database we used for training and testing the ANN. This bottleneck can be tackled by adding more data entries to the database once these become available. This would allow an improvement of the reliability of our algorithm for successfully reporting glycation probabilities.

Experimentally, approaches like nano high performance liquid chromatography/electrospray ionization/tandem mass spectrometry can be utilized to determine the ratio of glycated to total peptides ^45^. It should be kept in mind that usually amino acid glycation is not resulting in a “black or white” pattern (i. e., all peptides carrying the modification or none) but more like a gradual probability scale. Once more data on arginine glycation becomes available, we aim to present a tool based on our algorithm that analyzes a protein sequence provided by the user in FASTA format for the presence of arginine residues that are potentially glycated. The output would be given as an arginine glycation probability at a specific position of the protein in percent. This approach could allow narrowing down the amount of arginine residues that can preferentially become AGE-modified. Such a tool is envisioned to enable a more directed protein-AGE-arginine analysis saving time and funds for the researcher.

We want to stress that efforts have already been developed in this area from which the present research is inspired. Reddy *et al*. ^30^ developed a methodology based on support vector machines (SVM) with which they were able to classify glycated and non-glycated lysine residues using the structural properties of amino acid residues. For that work, they had a reference database containing a total of 538 glycated and non-glycated lysine residues, with which they were able to obtain an accuracy of 0.7562, 0 being totally inaccurate and 1 being totally accurate. Recently in Yu *et al*. ^26^ achieved a considerable improvement in the classification process of lysine glycations with SVM, working with a database of more than 6000 items, reaching a high accuracy of 0.88.

In comparison with the work presented here, it should be emphasized that although they are different methods (classification with SVM versus prediction with ANN), the highest precisions achieved are of similar magnitudes. However, there are several considerations to be stressed; first, the fact that our work had a very small base of only 54 peptides (both glycated and non-glycated peptides), which made the learning process of the neural network more complicated, and second the fact that what we performed in this project is an exact prediction of the probability of glycation, while the cited study and other similar studies on which this one is based ^25,27,29^ are founded on a classification between groups of peptides where there is glycation and where there is no glycation. It is important to note that all other studies prior to the one developed by Yu *et al*. ^26^ achieve, relatively speaking, lower accuracy.

Our algorithm shows that the most important characteristics determining arginine glycation probability are the sequence of amino acids, polarizability, amino acid mass, normalized van der Waals volume and hydropathy while torsion angle, hydrophobicity and isoelectric point seem to be of lesser importance (Fig. 4). When simultaneously considering two characteristics (two-case) polarizability and normalized van der Waals volume stand out as being most important for determining glycation probability (Fig. 5). The errors become lower when considering these two characteristics, showing that probably a combination of several factors predisposes an arginine residue for glycation. Scheck *et al*. ^21^ made the observations that polar residues like tyrosine (large van der Waals volume) and negatively charged ones seem to influence glycation probability. Certainly, it is possible that more than two properties of neighboring amino acids are relevant for the determination of arginine glycation probability. This question is planned to be addressed in future work.

## CONCLUSIONS

In conclusion, we herein present the conceptual framework that allows predicting the glycation susceptibility of arginine residues in peptides. Arginine modification by glycation is emerging to be highly relevant, perhaps even more so than lysine modification ^9,34^. Whereas several research groups addressed the question how to predict lysine modification, to our knowledge, we present the first attempt at predicting arginine glycation. At the same time, this study has been carried out using ANN on a very limited database. This is relevant given that previous studies on lysine have been carried out with the SVM method on databases of considerable size.

The present work focused on obtaining an accurate estimation of the probability of glycation in arginine. Promising results were obtained by taking combinations of two or three amino acid characteristics for such estimation. We identified that a combination of three characteristics (sequence of amino acids, amino acid mass and isoelectric point) gives the smallest mean absolute error (15.42). Combinations with other characteristics such as normalized van der Waals volume and hydropathy yield similar results. This key finding suggests that arginine glycation (and potentially glycation in general) is mostly influenced by the combination of these factors. Experimental approaches are needed to confirm this result.

Our work is aimed at the researcher who requires information on whether a certain arginine residue might be the target of reactive dicarbonyls and if so, to which extent. More than just reporting qualitative aspects we provide a strategy to receive quantitative information on the glycation probability of individual arginine residues. Therefore, the most probable “hits” would be the ones to whose experimental characterization would be applied preferentially. Overall, our approach is not only positioned to integrate into the landscape of previously published algorithms for the estimation of lysine residue glycation but to extend it in a meaningful way.

## ACKNOWLEDGEMENTS

Computational resources were supported by the biophysical systems laboratory of the Universidad Iberoamericana Puebla. Ulices Que-Salinas acknowledges the financial support provided by CONACyT México through grants: Estancias Posdoctorales Nacionales grant no. I1200/224/2021.

