## Supplementary material - SI and Figure S1 for "On the prediction of arginine glycation using artificial neural networks"

### 1.1 Table of properties

For the table of amino acid properties where authors used eight properties, amino acid sequence of the peptide (SoA), hydropathy (Hyd)[1], mass (Mas)[2], hydrophobicity (Hyp)[3], polarizability (Pol)[4], normalized van der Waals volume (vdW)[5], torsion angle (ToA)[6], and isoelectric point (IEP)[7], see Tab. S1.

To generate the SoA feature we arranged in descending order the amino acids by the hydropathy index. We set the 0 to the closest number (Glycine) and start an increasing/decreasing numbering using only integers. If they have the same value we order them by mass.

### 1.2 Table of vectors

For the table of created vectors see Tab. S2.

### 1.3 Artificial neural network operation (ANN)

To illustrate an ANN operation, consider the training process related to the MLP employed for the case considering hydropathy of the peptide's amino acids (case IB). For this we have 11 input neurons (one neuron for each amino acid), tree hidden layers with 4, 3 and 2 neurons, and only one output neurons for the estimation of the probability of glycation "G", where the relation between an input vector $\hat{P}$ (that represent all the patterns in the training set) and the $G_k$ output value is determined by the expression:

$$G_k = C\left(\tilde{b}_k + \sum_{l=1}^{20} \tilde{w}_{lk} \ * B\left(\tilde{b}_l + \sum_{j=1}^{40} \tilde{w}_{jl} * A\left(\sum_{i=1}^{11} w_{ij} * \hat{P} + b_j\right)\right)\right)$$

where $w_{ij}$, $\tilde{w}_{jl}$ and $\tilde{w}_{lk}$ are the connection weights between the neurons in input-hidden, hidden-hidden and hidden-output layers, respectively. The weights allow the perceptron to evaluate the relative importance of each of the outlets[8]. Neural network algorithms learn by discovering better and better weights that result in a more accurate prediction. There are several algorithms that are used to fine tune the weights, the most common is called backpropagation, which is used for all cases in this work. $b_j$, $\tilde{b}_l$ and $\tilde{b}_k$ are some extra weights called bias operating as thresholds. These weights allow you to move the curve of the activation function up and down or left and right on the number graph. This enables the numerical output of the perception to be adjusted. $A$, $B$ and $C$ are the ReLu activation functions between each consecutive layer.

To ensure that an ANN gives satisfactory results, its weights must be adjusted. The way to do this is by implementing a supervised training algorithm, that consists of introducing 38 examples into the ANN in which their training can be oriented in the right direction. Each example consists in *p* inputs denoted by $\hat{P}_{\mathrm{p}}$ and *k* outputs denoted by $\hat{G}_k{}^p$, where the index $p = 1$ ,…,38 represents the number of patterns used in the network training process. Thus, the task now is to find a way to minimize the error between the estimations $G_k{}^p$ made by the network (the outputs) and the corresponding values $\hat{G}_k{}^p$ expected to be predicted by the ANN during the training stage. This is done by a minimization of the weight dependent error function $E(\vec{w})$ (also called cost function)[9], which has several expressions in the literature. For this work, we consider one of the most recurrent expressions for the error function, defined for case I as

$$E(\vec{w}) = \frac{1}{38}\sum_{p=1}^{38}\sum_{k=1}^{1}\frac{1}{2}\left(G_k{}^p - \hat{G}_k{}^p\right)^2$$

The minimization of the error function is made by the iteration of the Adam algorithm, known as *backpropagation*[10,11], which searches the direction in which the error decreases the most on each epoch. One epoch means that each one of the 38 examples in the training sample of the data set has had an opportunity to update the internal model parameters (the weights). This search is done by changing the value for each $w_{ij}$ in the next epoch $t_e + 1$, propagating the error at epoch $t_e$ from output to input neurons by the rule

$$w_{ij}(t_e + 1) = w_{ij}(t_e) - \gamma \frac{\partial E(t_e, \vec{w})}{\partial w_{ij}(t_e)} + \alpha \triangle w_{ij}(t_e)$$

with $0 < \gamma < 1$ called the *learning rate*[12], $\alpha \triangle w_{ij}(t_e)$ is called the momentum term, added for preventing getting trapped in a local minimum, and $0 < \alpha < 1$ another parameter to be adjusted. The number of epochs for the training process is a hyperparameter that defines the number times that the learning algorithm will work through the entire training database. This whole process is carried out for the training of each of the cases studied, considering that due to the small size of the database, the results presented below for each case were averaged over the predictions made by 100 different ANNs already trained. It is worth mentioning that for all architectures of this study, we have used a rectified linear unit (ReLu) as an activation function for the layers[12-14].

The specific architecture of the ANNs used for each case is presented in Table [3]. Regarding the number of epochs, this value was adjusted for each case according to the time when, epoch by epoch, the improvement in accuracy was negligible compared to the number of epochs invested for such improvement. This is measured by considering the value of the cost function applied to the validation set at the end of each epoch, considering that the network has not been trained with it at all. This not only allows to stop the learning process when a certain desired accuracy has been reached, but also allows to monitor the process itself and to ensure that the ANN does not start to overtrain; that is, that the error does not start to increase for the validation set as it decreases for the training set. Finally, for each one of the study cases we have developed a database in order to train and test the ANN, selecting in general around 15% of the data of each database for the corresponding test set, and another 15% for the validation set. Considering the small number of proteins available to form the databases, a series of different distributions of the data sets was performed to ensure that the results were consistent as they were not conditioned to a specific distribution of the proteins in the training, validation, and prediction sets.

### 1.4 Results for case 2

The following figures shows the results of MAE and MAPE for the 28 property combinations of case 2.

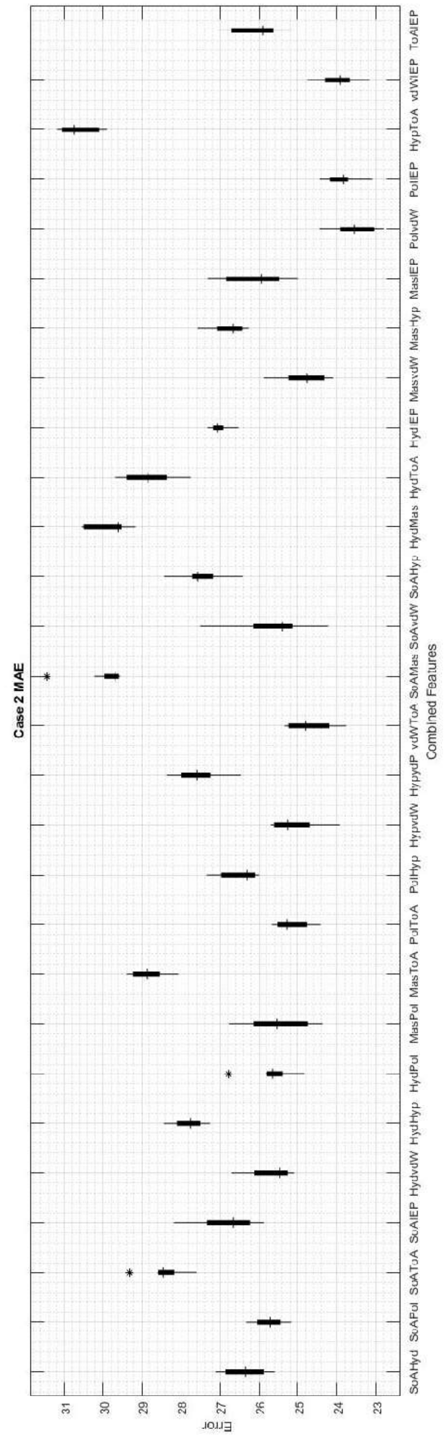


(a)

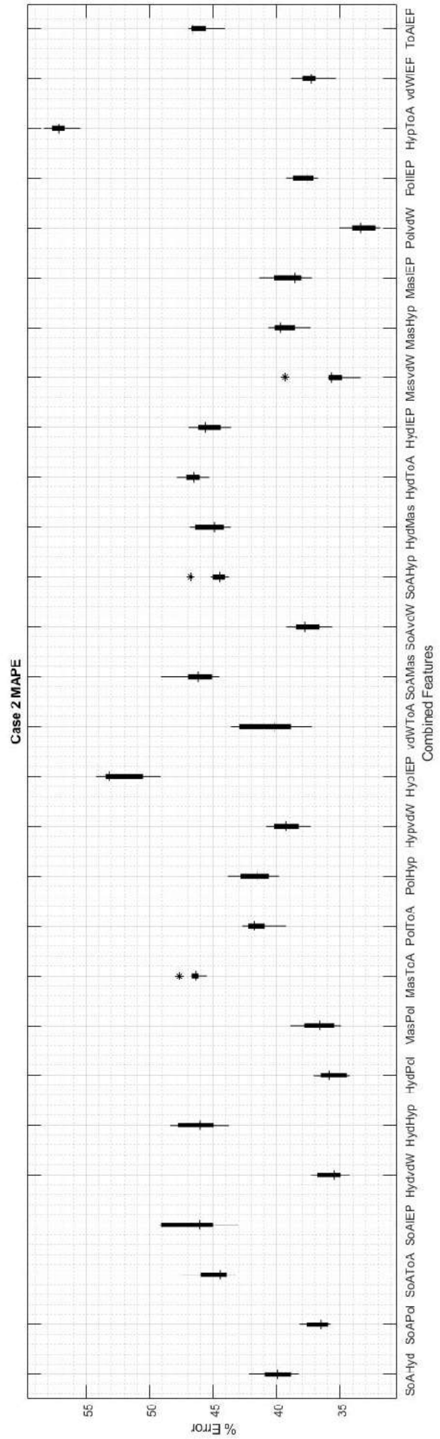


(b)

**Fig. S1.** Box plot of the results for case 2 for all the combinations of case 1, a) shows the values obtained for MAE, b) shows the values obtained for MAPE.
